# Gaudichaudione H inhibits Herpes Simplex Virus-1 replication by regulating cellular nuclear factor-κB in an interferon-γ-inde-pendent manner

**DOI:** 10.1101/2023.01.06.523065

**Authors:** Jiling Feng, Yuexun Tang, Wenwei Fu, Hongxi Xu

## Abstract

The highly prevalent herpes simplex virus type 1 (HSV-1) causes keratoconjunctivitis and encephalitis. Viral DNA polymerase-inhibiting nucleoside analogs (such as acyclovir) are standard treatment agents against HSV infections but are limited by severe drug resistance issues. Thus, new antiviral agents with novel targets are urgently needed. Earlier, we investigated the anti-cancer, anti-inflammatory, and antibacterial bioactivities of *Garcinia sp*. Here, we report that non-cytotoxic concentrations (< 500 nM) of Gaudichaudione H (GH, isolated from *Garcinia oligantha Merr*.) potently inhibits HSV-1 replication *in vitro* without affecting viral entry or attachment. GH inhibits the expression of the viral proteins ICP0, ICP4, and ICP27 without affecting their mRNA levels. In Vero cells, GH enhanced STAT1 and 3 phosphorylation, which occurs downstream to interferon (IFN)-γ activation during viral infections. However, pharmacological/genetic inhibition of IFN-γ failed to suppress the GH-mediated inhibition of HSV-1 replication, indicating that GH exerts antiviral effects independent of IFN. Further mechanistic studies suggest that GH inhibits HSV-1 replication, at least partially by inhibiting cellular NF-κB activation. Moreover, GH prolonged the survival rate of KOS-infected mice by 25% (n = 5). In conclusion, GH treatment inhibits HSV-1 replication both *in vitro* and *in vivo*; therefore, it can be developed as an antiviral.

**Importance:** Very few therapeutic drug options are available to treat herpes simplex virus-1/2 which cause myriad debilitating diseases. We screened eight *Garcinia* compounds and found Gaudichaudione H was the most effective compound at non-cytotoxic concentrations. Further mechanism study illustrates that GH inhibits HSV-1 replication, at least partially by inhibiting cellular NF-κB activation. Natural compound is a promising resource of new antiviral agents with different targets that has ability to treat resistant viral strains.

## 1. Introduction

Herpes simplex viruses (HSVs) comprise the HSV-1 and HSV-2 types, which primarily infect lips and external genitalia by forming vesicular dermatitis [1]. According to World Health Organization (WHO), around 67% (aged <50 and) and 13% (aged 15–49) people worldwide have HSV-1 and HSV-2 infection, respectively [2]. Upon infection, HSV is a lifelong companion that stays latent until host immunity is weakened. During this latency period, HSV usually hides in the trigeminal ganglia (TG) or dorsal root ganglia (DRG), and later travels to the central nervous system, resulting in serious diseases such as Alzheimer’s disease and encephalitis [3, 4].

Nucleoside analogs, incsluding acyclovir, penciclovir, and their derivatives, are the most effective antivirals in treating HSV infections [5]. However, resistance to these antivirals, which can develop within a week of treatment, poses a great challenge to the treatment of HSV infection using these drugs [6]. Nucleoside analog’s function as HSV DNA replication inhibitors by impeding viral thymidine kinases or viral DNA polymerases, and alterations in these enzymes confer drug resistance [7]. Viral helicase inhibitors and algae- or plant-derived compounds are being developed as second-line treatment options for acyclovir-resistant HSV infections [8, 9].

Interferons (IFNs) are the most important players of the host defense system [10]. Type I IFNs activate the Janus kinase (JAK)–signal transducer and activator of transcription (STAT) signaling pathway, which plays an important role in autoimmune protection. Activated JAKs phosphorylate and activate STATs, which allows them to dimerize, translocate to the nucleus, and regulate gene expression [11]. Nuclear factor (NF)-κB activation is another major antiviral innate response, which can be triggered by various signals induced by cellular receptors from different pathways. NF-κB signaling, which is activated by various stimuli such as viral or bacterial infection and exposure to proinflammatory cytokines and stress, is also an important intracellular antiviral pathway.

HSV1-induced persistent NF-κB nuclear translocation increases the virus replication efficiency [12]. Therefore, antiviral research mainly focuses on inhibiting nuclear translocation of NF-κB. Tumor necrosis factor alpha (TNF-α) or viral infection can cause targeted destruction of IκB and nuclear translocation of NF-κB. Following translocation, NF-κB mediates immune, inflammatory, or anti-apoptotic responses.

Traditional Chinese herbal medicines have long been used to prevent and treat viral infectious diseases in China and other oriental countries. They are an important source for discovering antiviral agents. Products from *Garcinia sp*., such as the well-known gamboge, have been traditionally used to treat sores and carbuncles. For years, we have focused on studying the bioactivities of compounds extracted from *Garcinia sp*. Here, we screened the following eight compounds derived from *Garcinia sp*. for their HSV replication inhibitory activity (compound information shown in Fig. S1A): i) Ypa (neobractatin) and ii) Ypb (isobractatin) from *Garcinia bracteate*; iii) OC (oblongifolin C), iv) GK (guttiferone K) and v) Garcinol from *Garcinia yunnanensis Hu*; vi) Gb-16 (gambogenic acid) from gamboge (*Garcinia cambogia*); vii) GH (Gaudichaudione H), and viii) A-4-f Oliganthin H from *G. oligantha Merr*. These eight compounds reportedly exert various bioactivities, such as inhibiting cancer cell proliferation, metastasis, and quiescent cancer cell reactivation, as well as antioxidant and anti-inflammatory activities [13-16]. Our screening revealed that GH is the most effective *Garcinia* compound in inhibiting viral replication at non-cytotoxic concentrations. Thus, we suggest that a natural compound from *Garcinia sp*. exerts antiviral activity and could potentially be developed as an inhibitor of HSV-1 replication.

## 2. Materials and Methods

### 2.1 Compound and chemicals

GH (with a purity greater than 98%) was extracted from the *Garcinia* oligantha Merr. (Clusiaceae) in our laboratory [17]. Ruxolitinib (11609, with purity ≥ 98%) was purchased from Cayman.

### 2.2 Cell line and viral strain

African green monkey kidney cells (Vero), human embryonic kidney cells (Hek293), human cervical cancer cells (HeLa), and mouse microglia cells (BV2) were purchased from ATCC, maintained in Dulbecco’s modified Eagle’s medium (DMEM) supplemented with 10% fetal bovine serum and a 1% antibiotic mixture (penicillin and streptomycin). The HSV-1 wild type strain KOS was provided by ATCC (VR-1493D). Green fluorescent protein (GFP)-fused KOS strain USGLP25 and ICP34.5 knockout strain KOS34.5 were previously constructed by Dr. Jia’s lab (see Acknowledgments).

### 2.3 Virus amplification

Vero cells were grown in monolayer until 80-90% confluent. Virus stocks were added at a MOI=1, and grow with cells for 72 h. Cells were frozen-thawed for 3 times and centrifuged. The supernatant containing virus was aliquoted, titrated, and kept at -80°C.

### 2.4 Cell viability assay

Cell viability was assessed by Thiazolyl Blue Tetrazolium Bromide (MTT). Cells (5 × 10^3^) were seeded in a 96-well plate and after treatment, 10 µL of MTT in phosphate-buffered saline (PBS) (5 mg/mL, M2003, Sigma) was added and absorbance was measured at 570 nm after 3 h incubation. Cell viability and IC50 values were determined using SPSS software 10.0.

### 2.5 Plaque assay

Vero cells were grown in 12-well plates in monolayer until 100% confluent and infected with the virus in DMEM without fetal bovine serum (FBS) for 1 h. The medium was then replaced with complete medium supplemented with 10% FBS, with or without indicated concentration of GH, and cells were incubated for 72 h. Next, cells were fixed with 4% paraformaldehyde and stained with crystal violet. Plaques were counted to calculate the virus titer.

### 2.6 Luciferase reporter assay

The NF-κB luciferase reporter plasmid is a kind gift from William Jia’s lab. A hundred thousand HEK293T cells were seeded in 24-well plates and incubated overnight. Cells were transfected with NF-κB Luciferase reporter plasmid using polyethylenimine (PEI, Sigma). The culture medium was refreshed 8 h post-transfection. Cells were kept in a humidified incubator for another 40 h, and the luciferase activity was determined using the dual luciferase assay system (E1500, Promega) according to the manufacturer’s protocols.

### 2.7 Virus titration assay

DNA was extracted from cell pellets using an EZNA Tissue DNA Kit (Omega Bio-Tek). Extracted DNA samples were subjected to quantitative polymerase chain reaction (qPCR) analysis using the SYBR Green Master Mix (Invitrogen) supplied with the primers for ICP27 and β-actin on a Quant Studio 6 Flex qPCR system (Applied Biosystems).

### 2.8 Real-time PCR

Total RNA was extracted using TRIzol reagent (Invitrogen) according to the manufacturer’s protocols. RT-PCR was performed with a one-step real time PCR using KAPA SYBR FAST One-Step qRT-PCR Universal (D-MARK Biosciences). Primers were used as previously described [18]. Results were calculated using 2 ^-ΔΔCt^ method.

### 2.9 Western blotting

Total protein was extracted using RIPA buffer (89901, thermo) and mixed with loading buffer (AM8546G, thermo), followed by heat block for 8 min at 99 °C. Protein samples were separated on SDS-PAGE (10% gel), electric-transferred to PVDF membranes, and incubated with primary antibody at 4°C overnight: beta-actin (4970), p-tat3(thy705, 9145), p-stat1(tyr701, 7649), p-eIF2a (Ser51, 9721), stat3 (4904), stat1 (9172) were from Cell Signaling Technology; NF-κB (ab16502), ICP0 (ab6513), ICP4 (ab6514) were from Abcam; ICP27 (sc-69806) were from Santa Cruz. Membranes were washed with TBST and incubated with secondary antibody (NEF812001EA, NEF822001EA, PerkinElmer) for 1 h. Membranes were visualized using a VersaDoc imaging system (Bio-Rad).

### 2.10 Confocal microscopy

P65 translocation were detected by confocal microscopy. Vero cells were grown on coverslips until 40-50% confluent, and infected with USGLP25 or KOS (MOI=0.1) with or without GH (0.5 µM) for 48 h. Cells were then fixed in 4% paraformaldehyde and permeabilized with Triton-X. Cells were blocked with 2% BSA for 1 h at room temperature. Cells were stained with primary and secondary antibody, and DAPI Fluoromount G (Electron Microscopy Sciences). Images were taken by a confocal microscope (Olympus).

### 2.11 Murine lethal challenge model

Procedure of animal study (ID01-QD026-2020v1.0) was approved by Ethics Committee of Nantong WuXi AppTec Animal Management and Use Committee. Briefly, 4-week-old female (weight, 17g) BALB/c mice were purchased from Shanghai Lingchang Biotechnology Co., Ltd. and kept under standard conditions. Mice were randomly divided into three groups (5 mice in each group): Mock, GH and pritelivir (positive control). Mice received intraperitoneal injection of GH (20 mg/kg/d) or pritelivir (3 mg/kg/d) or saline containing 10% Methylcellulose+5% dimethyl sulfoxide (DMSO, for mock group), once daily. After 0.5 h of the first drug administration, mice were anesthetized and total of 50 μL of virus suspension (3 × 10^6^ plaque-forming units [PFU], KOS) in ice-cold PBS was applied to the nares of mice. Animal behavior was monitored, body weight and mortality were recorded every day.

### 2.12 Statistics analysis

All data were presented as Mean ± SD of at least three independent experiments. Statistical analysis was performed using SPSS version 10.0. One-way ANOVA was used for multi-group comparison. * p < 0.05, ** p < 0.01, and*** p < 0.001 were considered statistically significant.

## 3. Results

### 3.1 GH poses antiviral effect against HSV-1 at a non-cytotoxic concentration

Antiviral screening should be performed at a non-cytotoxic concentration to avoid false positive results due to host cell death. So, we first determine cytotoxic concentration of the 8 *Gaicinia* compounds using MTT assay, and estimated the half cytotoxic concentrations (CC_50_) (Fig. S1B). The dosages showing >90% viability of the cells (framed by red boxes in Fig. S1C) were selected as the maximum dosages for screening antiviral activity. Next, we screened for antiviral activity using GFP-fused HSV-1 called USGLP25. Viral infection was quantified by monitoring changes in fluorescent signals in the infected Vero cells. Among the eight Garcinia compounds, GH showed the best antiviral activity. Cell viability assay showed that cytotoxicity of GH did not increase with virus infection (Fig. 1A, B), and fluorescence of USGLP25-infected Vero cells is shown in Fig. 1C. Subsequently, we performed plaque assay to confirm the antiviral activity of GH. Plaque assay illustrated that GH treatment significantly inhibited the virus titer of both USGLP25 and KOS in a dose-dependent manner (Fig. 1D, E). All these results suggest that GH treatment inhibits virus replication without causing additional cytotoxicity in Vero cells.

**Figure 1.**
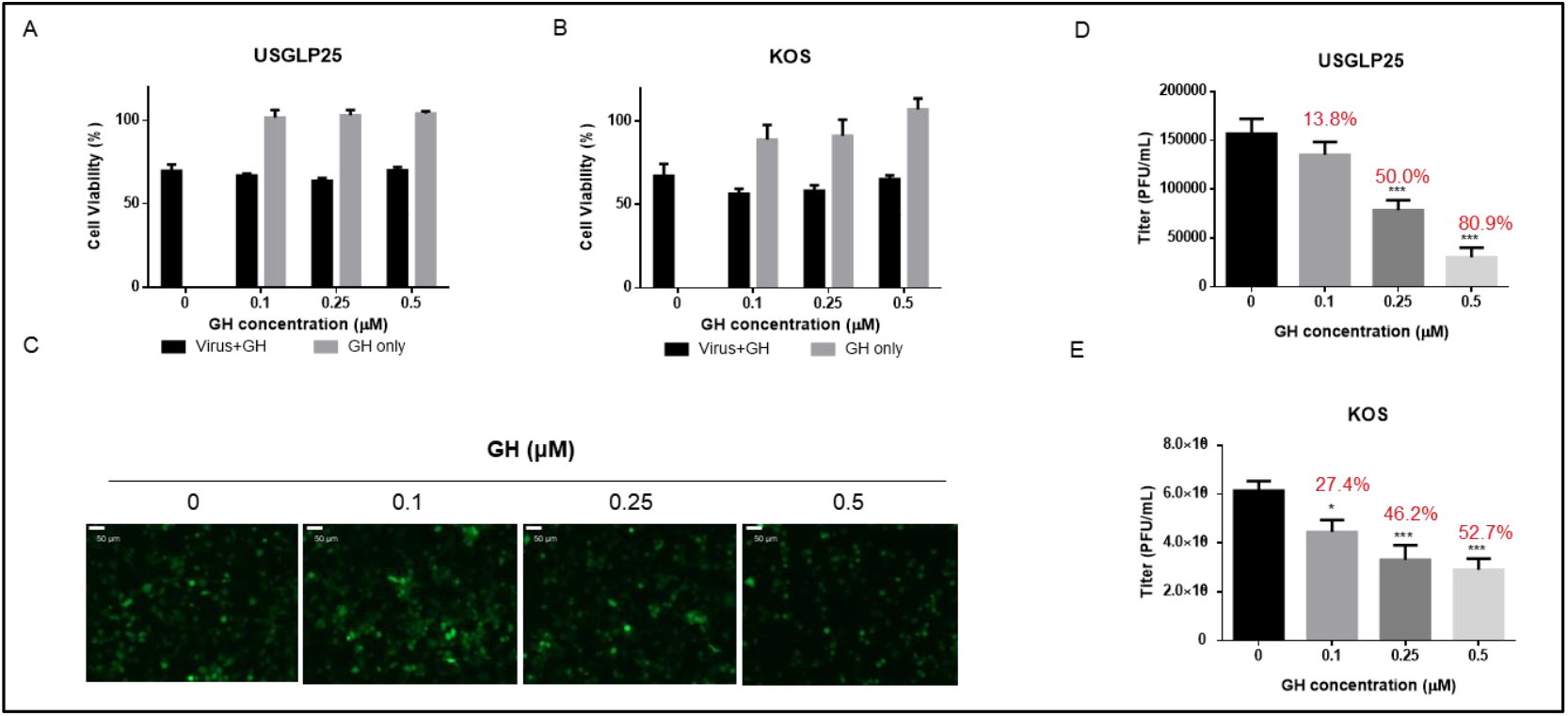
GH inhibits HSV-1 replication at a non-cytotoxic concentration. Vero cells were infected with USGLP25 (A, C, D, MOI=1) or KOS (B, E, MOI=1) with or without GH (0.5 μM) for 48 h. Cell viability was assessed using MTT assay (A, B) and virus titer was assessed by titration assay (D, E). Inhibition rates were calculated and shown in red . * p<0.05, *** p<0.001 compared to vehicle group. Immunofluorescence of USGLP25 infected Vero cells (C). Scale bar= 50 µm.

### 3.2 GH inhibits HSV-1 replication without affecting viral entry or attachment

To identify the viral infection step inhibited by GH, we performed a pre- and post-treatment assay. We treated Vero cells with GH at indicated concentrations either before or after infection with *HSV* viral strains and calculated the virus titer at 16 h post-infection. At concentrations > 0.1 µM, GH treatment significantly reduced the viral titers in pre- and post-treated batches. Since the inhibition observed was similar in pre- and post-GH-treated batches, it suggested that GH does not affect virus entry (Fig. 2A, B). To further reveal the inhibitory effect of GH on viral amplification, we performed a one-step viral growth assay. Using qPCR and ICP27 primers to quantify the viral replication, we studied the effect of GH on viral replication by simultaneously adding the virus and GH and calculating virus titers at 1 h and 16 h post-infection. ICP27 amplification was significantly reduced with GH treatment; this GH-mediated effect was observable only at 16 h but not 1 h post-infection. Thus, GH inhibits HSV replication without affecting viral attachment to host cells. Taken together, GH inhibits HSV-1 virus replication without affecting viral entry or attachment.

**Figure 2.**
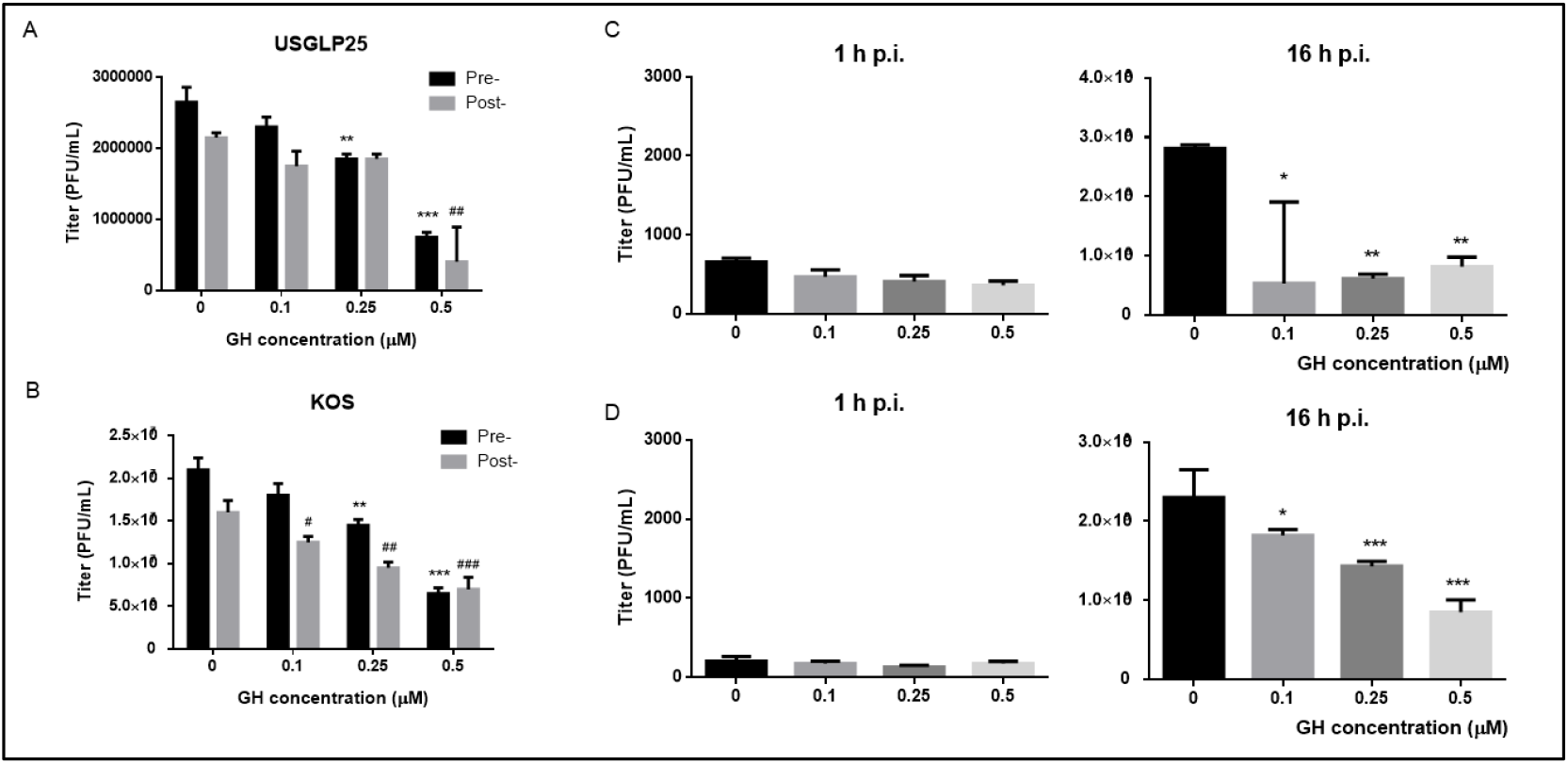
GH inhibits HSV-1 virus replication without affecting viral entry or attachment. (A-B) Vero cells were pre-treated with GH at the indicated concentration for 1 h, then infected with USGLP25 or KOS (MOI = 1); or Vero cells were infected with USGLP25 or KOS virus for 1 h and then incubated in medium containing the indicated concentration of GH. Virus titers were determined via plaque assay; ** p < 0.01, *** p < 0.001 compared to the vehicle group of pre-treatments; # p < 0.05, ## p < 0.01, ### p < 0.001 compared to the vehicle group of post-treatment. (C-D) Vero cells were infected with USGLP25 (C, MOI = 1) or KOS (D, MOI = 1) with indicated concentrations of GH for 1 h, and the medium was changed, containing the same concentration of GH without the virus. Cells were collected immediately (1 h post-infection) or after another 15 h of incubation (16 h post-infection) and stored at -80 °C. Virus titer was determined using qPCR assay with ICP27 primers; * p < 0.05, ** p < 0.01, *** p < 0.001 compared to the vehicle group.

**Figure 3.**
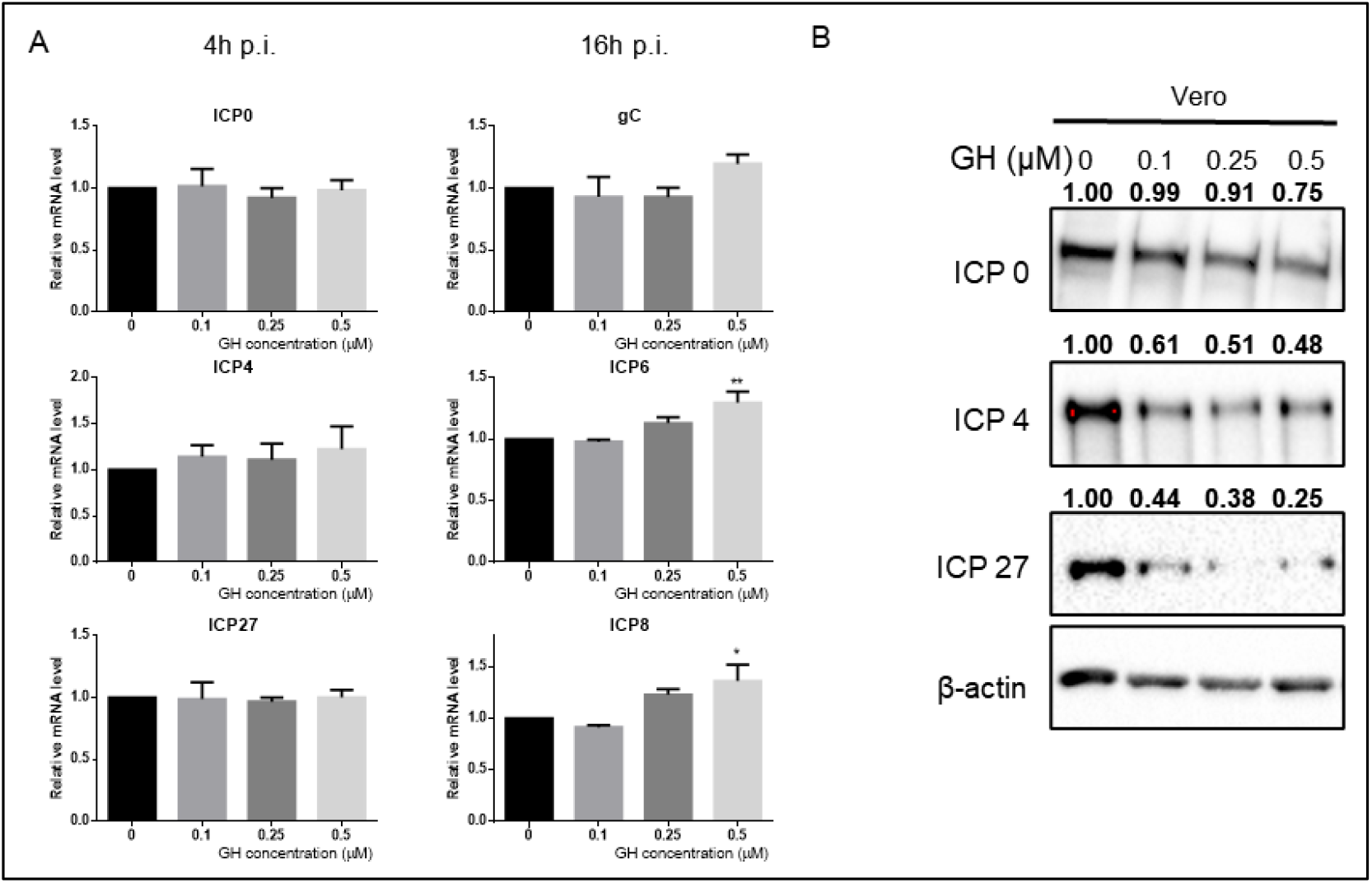
GH inhibits important viral genes at the protein level but not at the mRNA level. Vero cells were infected with KOS (MOI = 3) with the indicated concentration of GH (0-0.5 μM), (A) total RNA was extracted 4 h (ICP0, ICP4, and ICP27) or 16 h (ICP6, ICP8, and glycoprotein C) after virus infection. (B) total proteins were extracted at 5 h post-infection. β-actin served as the loading control. Quantification of band intensities relative to vehicle group was analyzed using Image J software and is shown above the bands.

Next, using qPCR, we checked mRNA levels of several important viral genes, including immediate early genes (ICP0, ICP4, UL54), early genes (UL29, UL39), and late gene (UL44). GH treatment did not affect transcription of any of these genes (Fig.3A). However, it significantly inhibited translation of the immediate early genes; protein levels of ICP0, ICP4, and ICP27 were decreased upon GH treatment (Fig.3B). These results suggested that GH serves to degrade the early viral gene-encoded proteins via host-cell antiviral mechanisms.

### 3.3 GH inhibits HSV-1 replication in an interferon-γ-independent manner

Next, we studied the effect of GH treatment on IFN pathways, the most common antiviral mechanism employed by the host immune system. IFN functions through JAK/STAT signaling resulting in activation of several kinases, including the double-stranded RNA-activated protein kinase (PKR). Once activated, PKR autophosphorylates and catalyzes phosphorylation of eukaryotic translation initiation factor 2 subunit alpha (EIF2α) [19]. In both KOS- and USGLP25-infected Vero cells, GH treatment increased p-STAT1, p-STAT3, and p-EIF2α expression, without affecting that of STAT1 or 3 (Fig. 4A). Next, we used STAT inhibitor to rescue the antiviral effect of GH treatment; we used ruxolitinib as a STAT/JAK inhibitor, which is an FDA-approved drug of myelofibrosis [20]. Ruxolitinib treatment inhibited p-STAT3 and p-STAT1 expression in KOS-infected Vero cells (Fig. 4B). However, virus titration assay revealed that ruxolitinib treatment did not significantly alter the antiviral effect of GH treatment in Vero cells (Fig. 4C).

**Figure 4.**
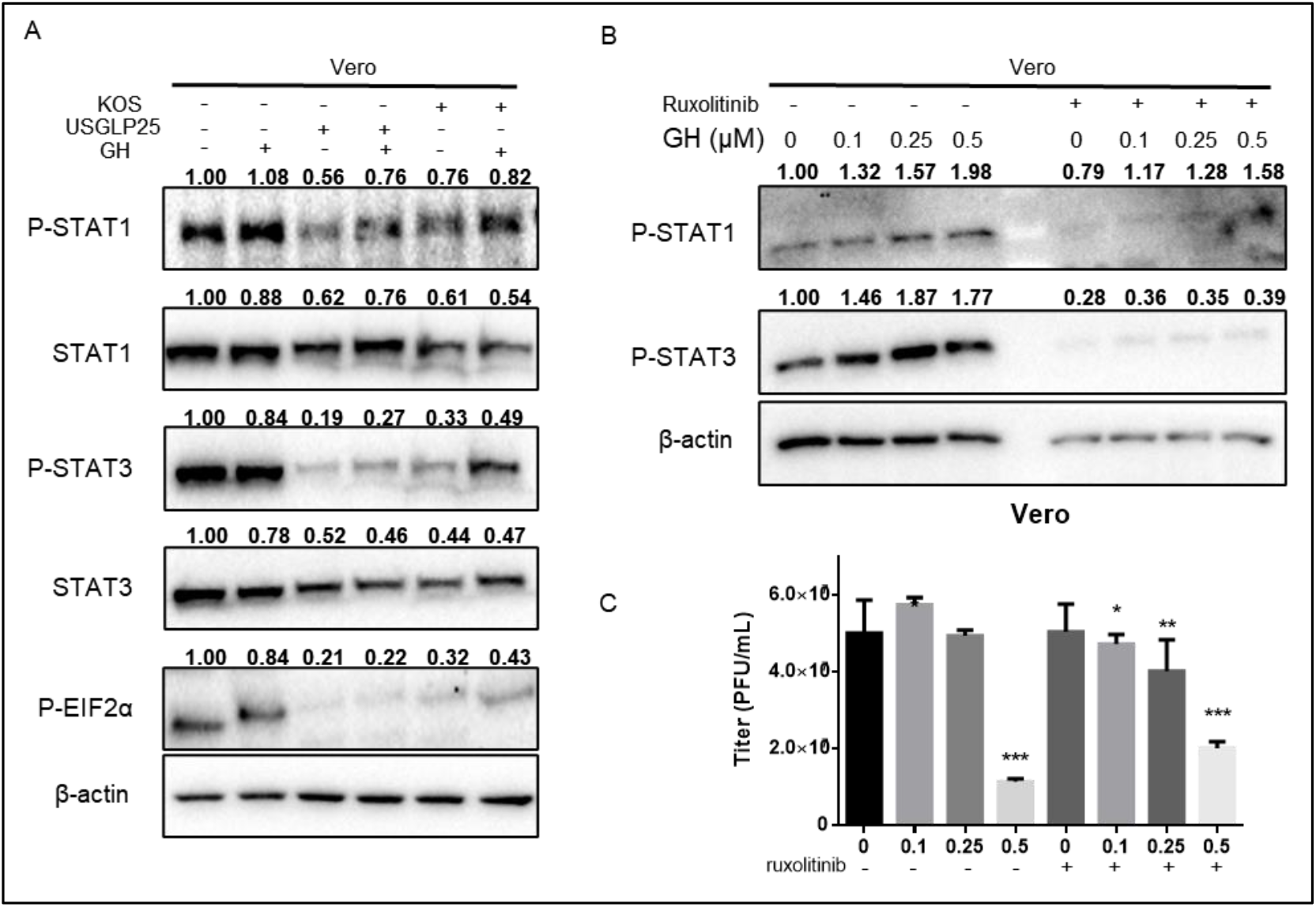
GH inhibits HSV replication via interferon γ-independent pathways. (A) Vero cells were infected with USGLP25 or KOS (MOI = 3) with or without GH (0.5 μM), and total proteins were extracted 16 h post-infection for the WB assay. (B) Vero cells were pretreated with ruxolitinib (125 nM) for 2 h and then infected with KOS (MOI = 3) with or without GH at indicated concentrations. Total proteins were extracted 16 h post-infection for the WB assay. β-actin served as the loading control. Quantification of band intensities relative to that of the vehicle group was analyzed using Image J software and is shown above the bands. (C) Vero cells were pre-treated with ruxolitinib (125 nM) for 2 h and then infected with KOS (MOI = 1) with or without GH at the indicated concentration. Virus titration was assessed by qPCR 16 h post-infection; * p < 0.05, ** p < 0.01, *** p < 0.001 compared to the vehicle group.

Vero cells have deficiency in IFN system due to the lack of IFN-alpha and IFN-beta genes, which explains the low levels of STAT phosphorylation observed in Vero cells upon virus infection [21]. Therefore, we next checked the effect of GH treatment on BV2 cells, a microglial cell line derived from C57/BL6 mice, which have strong IFN signaling. We found GH treatment enhanced STAT phosphorylation in uninfected BV2 cells but irrespectively inhibited STAT phosphorylation in KOS-infected cells (Fig. 5A). We predicted that treating BV2 cells with ruxolitinib would enhance the inhibitory effect of GH treatment on viral replication. Western blotting results showed that GH could no longer inhibit STAT1 and STAT3 phosphorylation with ruxolitinib (Fig. 5B). However, the GH inhibitory effect on virus replication in BV2 cells remained unchanged, irrespective of ruxolitinib pre-treatment (Fig. 5C). We repeated these experiments with KOS-infected human cervical cancer (HeLa) and human embryonic kidney (Hek293) cell lines; GH treatment inhibited STAT phosphorylation in both these cell lines (Fig. S2). Thus, GH treatment seemed to inhibit STAT phosphorylation only in cells capable of IFN production, indicating that GH does not exert its antivirus activity by activating IFN pathways.

**Figure 5.**
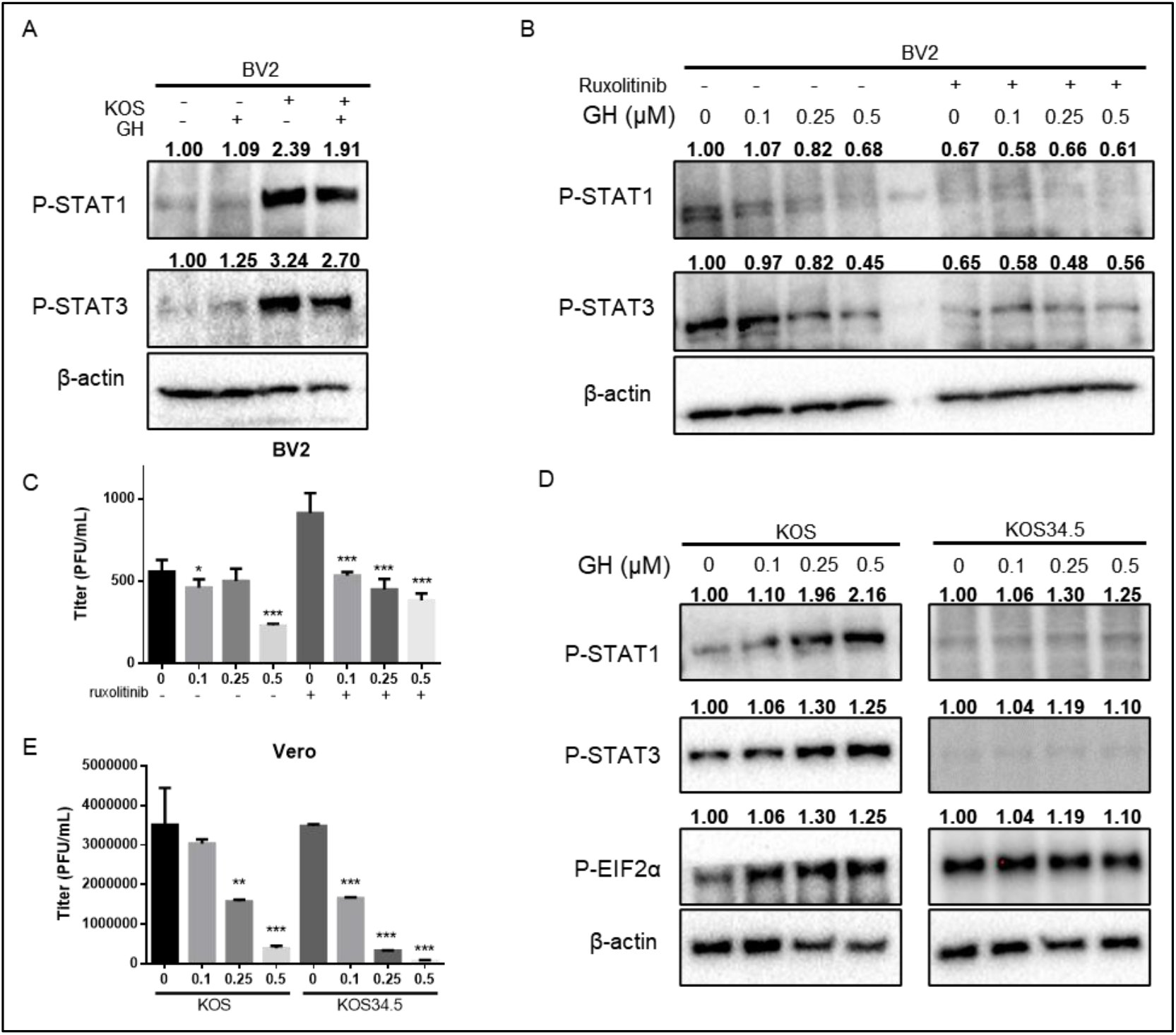
GH inhibits HSV replication in an interferon γ-independent manner. (A) BV2 cells were infected with KOS (MOI = 3) with or without GH (0.5 μM), and total proteins were extracted 16 h post-infection for the WB assay. (B) BV2 cells were pre-treated with ruxolitinib (125 nM) for 2 h and then infected with KOS (MOI = 3) with or without GH at indicated concentrations. Total protein was extracted 16 h post-infection for the WB assay. β-actin served as the loading control. (C) BV2 cells were pre-treated with ruxolitinib (125 nM) for 2 h and then infected with KOS (MOI = 1) with or without GH. Virus titration was assessed by qPCR 16 h post-infection. (D) Vero cells were infected with KOS or KOS34.5 (MOI = 1), with indicated concentrations of GH. Virus titration was assessed by qPCR 16 h post-infection. (E) Vero cells were infected with KOS or KOS34.5 (MOI = 3), with indicated concentrations of GH. Total protein was extracted 16 h post-infection for WB assay. β-actin served as the loading control. Quantification of band intensities relative to vehicle group was analyzed using Image J software and is shown above the bands.

ICP34.5 is implicated in many aspects of viral pathogenesis. It is targeted by host IFN responses to impede protein synthesis by counteracting PKR activity and reversing EIF2α phosphorylation [22]. We used a stable ICP34.5 knockout KOS virus called KOS34.5. Due to its limited infection activity in BV2 cells, we only performed KOS34.5 infection on Vero cells. Western blotting showed that GH treatment could no longer activate p-STAT1 and p-STAT3 in KOS34.5-infected Vero cells (Fig. 5D). However, it could significantly inhibit KOS34.5 replication in infected Vero cells (Fig. 5E). Taken together, we confirmed that GH treatment inhibits HSV-1 replication in an IFN-independent way. Next, we studied GH effect on other cellular pathways implicated in antiviral activity to find out how GH treatment inhibits HSV-1 replication.

### 3.4 GH inhibit HSV-1 replication partially by inhibiting NF-κB nuclear translocation

Transcription factors of the NF-κB/Rel family are important to inflammatory and immune responses. p65, a major functional subunit of NF-κB, translocates to the nucleus in HSV-1-infected cells [23]. We performed a luciferase reporter assay to check whether GH treatment inhibits p65 nuclear translocation. Indeed, GH treatment partially reversed the viral infection-induced p65 nuclear translocation (Fig. 6A). We also performed immunofluorescence assay to detect localization of p65. Both DMSO- and GH-treated uninfected cells exhibited a significant separation of p65 (red) and DAPI (blue), indicating that GH treatment did not significantly affect p65 nuclear translocation in uninfected cells (Fig. 6b). In KOS-infected cells, p65 was well colocalized with the nucleus, suggesting that p65 translocated into the nucleus upon virus infection (Fig. 6B-C). GH treatment significantly reduced (the pink color in KOS-infected cells and white color in USGLP25-infected cells) the overlap ratio of p65 and nucleus, indicating that GH treatment blocks viral infection-induced p65 nuclear translocation (Fig. 6B-D). Therefore, GH treatment may exert its antiviral effect by blocking p65 nuclear translocation.

**Figure 6.**
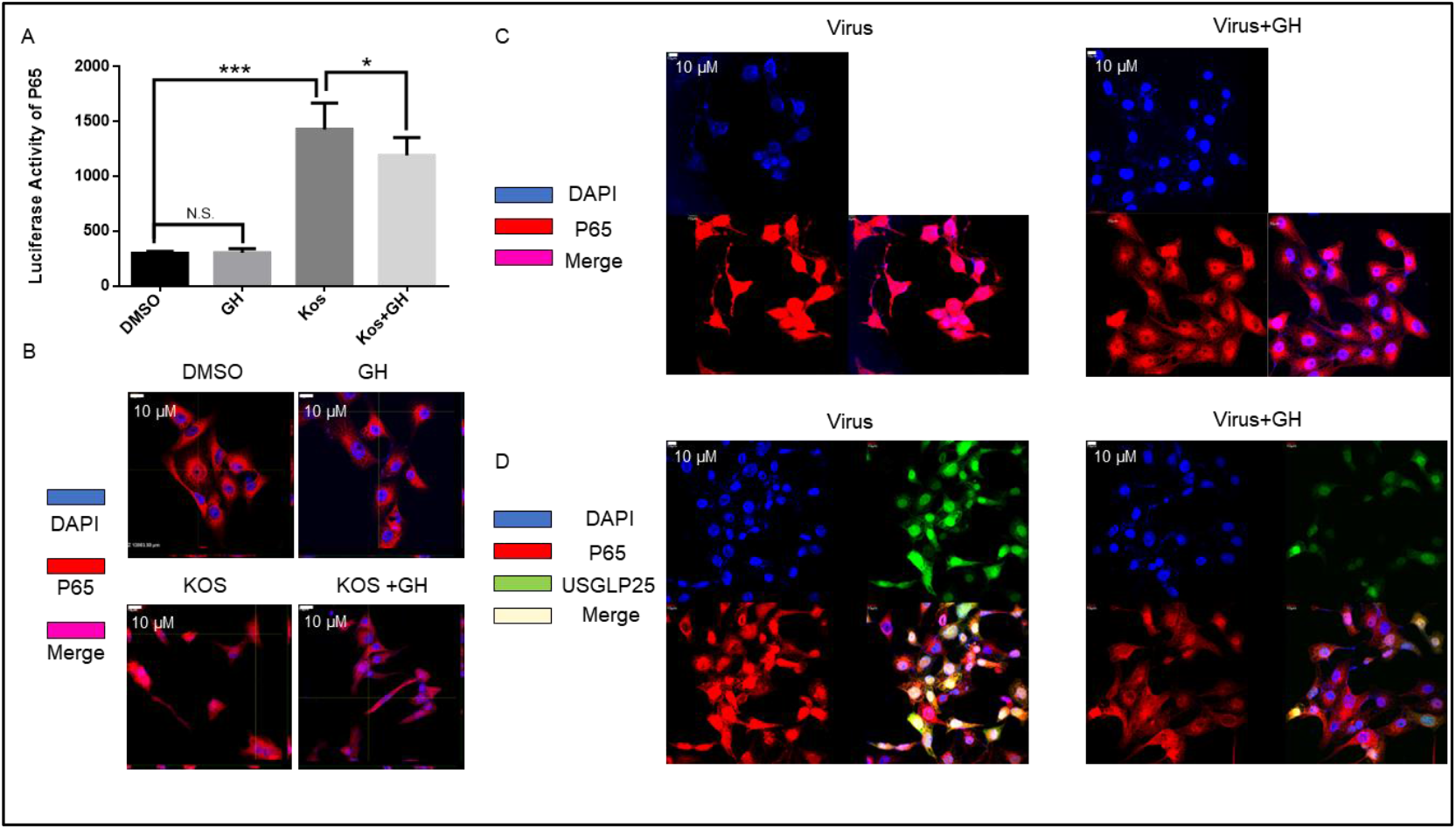
GH inhibit HSV-1 replication partially by inhibiting NF-κB nuclear translocation. **(A)** Vero cells were transfected with NF-κB luciferase reporter plasmid and infected with KOS (MOI=1), with or without GH (0.5 μM). Luciferase activity was determined after 48 h. Vero cells were infected with KOS (B, C, MOI=1) or USGLP25 (D, MOI=1), with or without GH (0.5 μM) for 24 h. Scale bar= 10 μm. Overlapping area showing pink (the colocalization of DAPI (blue) and p65 (red) for KOS, C) or white color (the colocalization of DAPI (blue), p65 (red), and USGLP25 (green), D) indicated nuclear translocation of p65. C-D: virus infected without (left lane) or with (right lane) GH.

### 3.5 GH inhibits HSV-1 replication *in vivo*

To confirm the in vivo antiviral effect of GH, we constructed a lethal-challenge animal model. Mice were randomly divided into three groups: Mock, GH, and pritelivir (positive control). GH and pritelivir were intraperitoneally administrated once daily. After administrating the first dose of the drug, mice were intranasally infected with KOS (3 × 10^8^ PFU). This infection protocol can result in a mortality rate of 90 to 100% within 6 to 10 days due to dissemination of disease [24]. On the third day after infection, the viability of the mock group was 40% (two of five mice survived), whereas that of the GH group was 100% (all five survived). On the fourth day, viability of the mock group was 0% (all five mice died), whereas that of the GH group was 40% (two of five mice survived). The remaining two mice died on the fifth day. Viability of the pritelivir group was 100% until day 5 (all five mice survived without obvious pathological changes) (Fig. 7A). Body weight of the mice was recorded every day (Fig. 7B). Thus, these results indicate that GH has antiviral activity *in vivo*.

**Figure 7.**
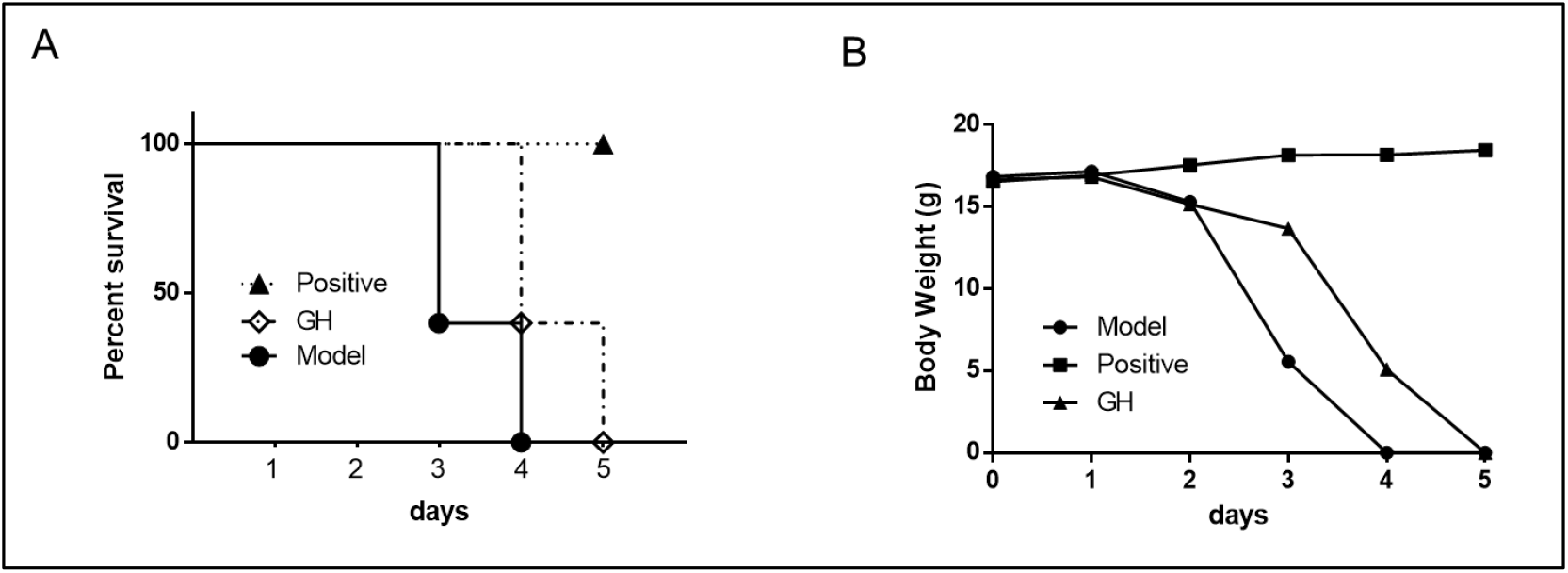
GH inhibits KOS replication in vivo using a lethal challenge model. Mice were infected intranasally with KOS and treated once daily with GH (20 mg/kg/d) or pritelivir (positive, 3 mg/kg/d), or vehicle (saline containing 10% Methylcellulose+5% DMSO) as indicated for 5 days beginning 0.5 h before virus infection. Survival rate(A) and change in body weight(B) were recorded every day. Five animals from each group were used.

## 4. Discussion

Results presented here demonstrate that GH, a *Garcinia* compound, can inhibit HSV-1 replication both in vitro and in vivo, in an IFN-independent way. GH acts partially through nuclear translocation of NF-κB. NF-κB is a critical regulator of the host innate immune system. However, HSV-1 transiently activates NF-κB to initiate its early infection and subsequently escapes the host immune surveillance using its gene products [25]. NF-κB is necessary for prevention of host cell apoptosis during the early stage of HSV infection. Nevertheless, it is still widely accepted that NF-κB has antiviral effects. Many human viruses, such as hepatitis C virus (HCV), human immunodeficiency virus (HIV), Kaposi’s sarcoma-associated herpesvirus (KSHV), and Epstein–Barr virus (EBV), have evolved to harness the host cellular NF-κB pathway [26]. However, several approved NF-κB-targeting drugs have antiviral effects. For instance: 1) thalidomide, a US Food and Drug Administration (FDA)-approved immunomodulatory agent, inhibits HSV and varicella zoster virus coinfection in cancer patients as well as pseudotumoral recto-sigmoid HSV-2 coinfection in HIV patients [27]; and 2) acetylcysteine, a reactive oxygen species scavenger, is used as a mucolytic agent. Acetylcysteine exerts antiviral effect against HSV-1 and HSV-2 [28]. Given the role of the host cellular NF-κB signaling in HSV viral replication, modulation of this pathway seems to be an alternative approach to prevent viral infections. Here, we shed light on the possibility that GH exerts antiviral effect by regulating NF-κB. GH treatment suppresses HSV-1-stimulated NF-κB pathway activation and p65 nuclear translocation. Further studies verifying the effect of GH treatment on p65 protein and its phosphorylation levels are needed to define the exact effect of GH treatment on p65. Effectiveness of p65 translocation inhibitors in rescuing the in vitro and in vivo effects of GH treatment must also be studied.

IFN, the most widely studied aspect of the innate immune system, is the first line of defense against invading pathogens [29]. Recently, focus is shifting to IFN-independent anticancer and antiviral mechanisms employed by cells. Mammalian stimulator of interferon genes (STING) possesses widespread IFN-independent activities that can restrict HSV-1 and aid tumor immune evasion [30]. STING-mediated death in T cells could help tumors evade immune control. Furthermore, in triple-negative breast cancers, ligand-dependent corepressor (LCOR) binds IFN-stimulated response elements (ISREs) driving immune escape and immune-checkpoint blockade in an IFN-independent manner [31]. We demonstrated that GH treatment exerts antiviral effect in an IFN-independent manner. Future studies can explore the effect of GH treatment on STING- and LCOR-related pathways.

Nucleoside analogs such as acyclovir, valaciclovir, penciclovir, ganciclovir, and famciclovir are the most well-known first-line antivirals used against HSV. Here, we used FDA-approved pritelivir as the positive control for *in vivo* tests, which has not been approved by the China FDA (CFDA). The following drugs are CFDA approved antivirals—Recombinant Human Interferon A2A Cream for facial and genital herpes caused by HSV-1/2 infection; Recombinant Human Interferon α2b Gel for zoster, labial and genital herpes; and Viru-Merz (Tromantadine Gel) to treat primary and recurrent HSV. Besides, licensed vaccines against HSV are currently lacking, but efforts are being made to develop vaccines against varicella-zoster virus, which has similar structure to HSV [32]. The HSV vaccine candidate from GlaxoSmithKline seems to be the most promising among several vaccine candidates being studied clinically [33]. Effective antiviral agents among natural products that inhibit viral infection while causing minimal damage to host cells would be promising.

Acyclovir resistance, which causes recurrence of symptoms and increases risk of complications, is still the main challenge in antiviral therapy. Since most antiviral drugs target viral DNA polymerases or viral thymidine kinases (TK), effective options to treat drugresistant HSV are nil [34]. Here, USGLP25 has impaired viral TK gene expression. Therefore, it can be considered an acyclovir-resistant HSV strain. Given that GH treatment significantly inhibits USGLP25, GH might be effective against drug-resistant HSV. Future studies must verify whether GH can inhibit the replication of acyclovir-resistant viruses and act synergistically with acyclovir in inhibiting viral replication. Our study suggests that compounds from *Garcinia sp*. can be potentially developed as antiviral drugs.

## Author Contributions

Supervision and funding acquisition, H-X.X.; Project administration and manuscript writing, J-L.F.; Data curation, Y-X.T.; Statistic analysis and software, W-W.F. All authors have read and agreed to the published version of the manuscript.

## Funding

This research was funded by the Key-Area Research and Development Program of Guangdong Province (No. 2020B1111110003); the NSFC-Joint Foundation of Yunnan Province (No. U1902213); National Natural Science Foundation of China (No. 82104209).

## Institutional Review Board Statement

Procedure of animal study (ID01-QD026-2020v1.0) was approved by Ethics Committee of Nantong WuXi AppTec Animal Management and Use Committee (2020-11-19).

## Acknowledgments

We thank Professor William Jia and Dr. Judy Ding (previously at UBC) for their guidance on the project and providing lab supplies.

## Conflicts of Interest

The authors declare no conflict of interest.

---

**Figure S1.**
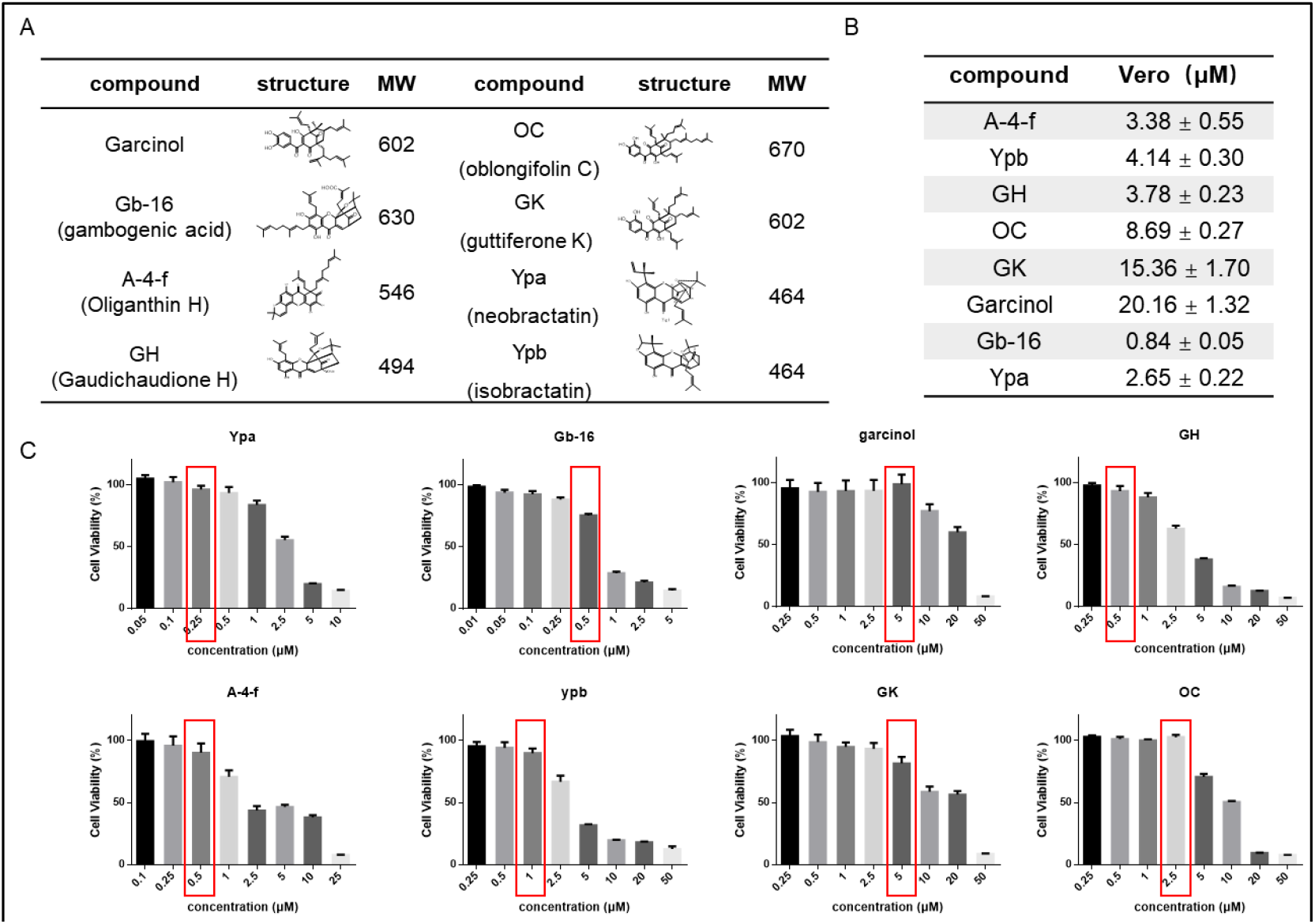
Compound information and CC_50_ of 8 Garcinia Compounds. (A) structure and molecular weight of 8 Garcinia compounds. (B) CC_50_ of the 8 compounds in Vero and HFF cell line. Cells were seeded in 96 well plate and treated with indicated concentration of compounds. Cell viability was assessed by MTT assay. CC_50_ was calculated using SPSS 10.0 software. (C) MTT assay of 8 compounds on Vero cells and the red boxes indicated the concentration of 90% viability.

**Figure S2.**
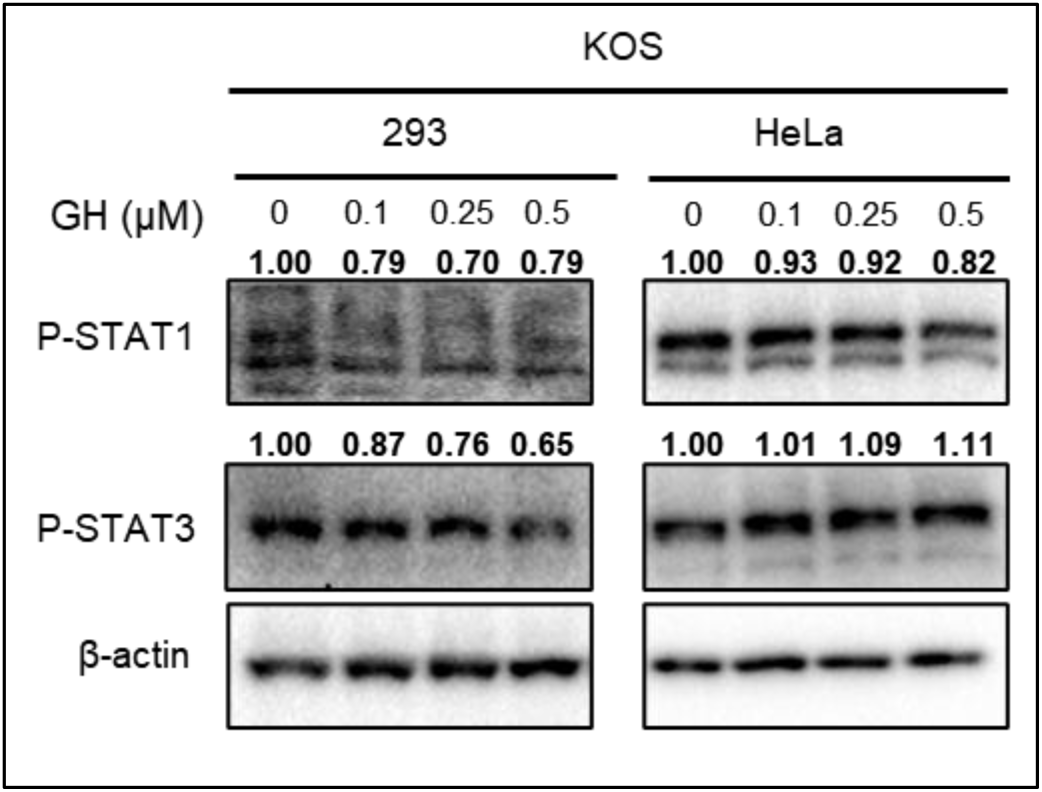
GH inhibited STAT phosphorylation in Hek293 and Hela cells. Hek293 and Hela cells were infected with KOS (MOI=3), with indicated concentrations of GH. Total protein was extracted 16 h post infection for WB assay. Beta-actin served as loading control. Quantification of band intensities relative to vehicle group was analyzed using Image J software and were shown above the bands.

